# Activity of Sotrovimab against BQ.1.1 and XBB.1 Omicron sublineages in a hamster model

**DOI:** 10.1101/2023.01.04.522629

**Authors:** Jean-Sélim Driouich, Ornéllie Bernadin, Franck Touret, Xavier de Lamballerie, Antoine Nougairède

**Author notes:** Corresponding author: Antoine Nougairède.

## Abstract

The successive emergence of SARS-CoV-2 Omicron variants has completely changed the modalities of use of therapeutic monoclonal antibodies. Recent in vitro studies indicated that only Sotrovimab has maintained partial activity against BQ.1.1 and XBB.1. In the present study, we used the hamster model to determine whether Sotrovimab retains antiviral activity against these Omicron variants in vivo. Our results show that at exposures consistent with those observed in humans, Sotrovimab remains active against BQ.1.1 and XBB.1, although for BQ.1.1 the efficacy is lower than that observed against the first globally dominant Omicron sublineages BA.1 and BA.2.

## Main text

Therapeutic monoclonal antibodies (mAbs) targeting the SARS-CoV-2 spike protein have been widely used during the current COVID-19 pandemic, particularly in immunocompromised patients in whom vaccination induces an inadequate immune response. However, almost all clinically approved mAbs have lost part or all of their neutralizing activity against the different sub-lineages of the Omicron variant that have successively spread globally and contain several mutations in the spike protein associated with potential escape from humoral immunity and higher transmissibility (Campbell et al., 2021; Cao et al., 2022; Cox et al., 2022). While Bebtelovimav, Cilgavimab, Imdevimab and Sotrovimab retained some activity against BA.5, only Sotrovimab maintains partial activity against BQ.1.1 and XBB.1, BA.5 and BA.2 respective subvariants with increasing incidence in the USA and Europe (Arora et al., 2023; Imai et al., 2022; Planas et al., 2022; Touret et al., 2022). It has been shown that despite its partial loss of in vitro neutralising activity against BA.2 and BA.5, Sotrovimab exhibits antiviral activity in Syrian hamsters against these Omicron variants (Park et al., 2022; Uraki et al., 2022). It is therefore urgent to determine whether this mAb retains activity in vivo against BQ.1.1 and XBB.1 at exposures similar to those observed in humans.

In this study, the efficacy of Sotrovimab against four clinical strains of Omicron variant (BA.1, BA.2, BQ.1.1 and XBB.1) was assessed in a hamster model using an ancestral B.1 strain as reference. Three days before the intranasal infection, groups of animals received pre-exposure prophylaxis by intramuscular injection of increasing doses of Sotrovimab and were compared with control treated animals receiving an isotype control mAb (Palivizumab)(Fig 1A). Overall, results showed that Sotrovimab exhibited an antiviral activity against all viruses, although less marked for BQ.1.1, in agreement with recently published in vitro data based on a VeroE6/TMPRSS2 cell assay with replicating virus (Touret et al., 2022). Indeed, administration of Sotrovimab always dose-dependently reduced infectious titers in lungs. When compared with corresponding control treated animals, mean reductions ranged between 82.33 and 99.99% for the ancestral strain, between 84.58 and 99.78% for BA.1, between 96.47 and 99.86% for BA.2, between 78.42% and 98.28% for BQ.1.1 and between 93.62% and 99.74% for XBB.1 (Fig 1B-C). This decrease was significant for all doses of Sotrovimab used against all viruses (*p* values ranging between <0.0001 and 0.0369), except with the dose of 2mg/kg for the BQ.1.1. Furthermore, administration of Sotrovimab led to a reduction of viral RNA yields in nasal washes for all doses with all viruses (*p* ranged between <0.0001 and 0.0227), except with the dose of 0.7mg/kg for the ancestral strain and the BA.1 variant, and the dose of 7mg/kg for the BQ.1.1 (Fig 1D). In addition, clinical monitoring of animals showed at 3 days post-infection (dpi) significantly higher normalized weight with the dose of 7mg/kg for the B.1 strain, the dose of 2 and 14mg/kg for the BA.2 variant and the dose of 2, 7 and 14mg/kg for the BQ.1.1 variant when compared with corresponding control treated groups (Fig S1).

**Figure 1:**
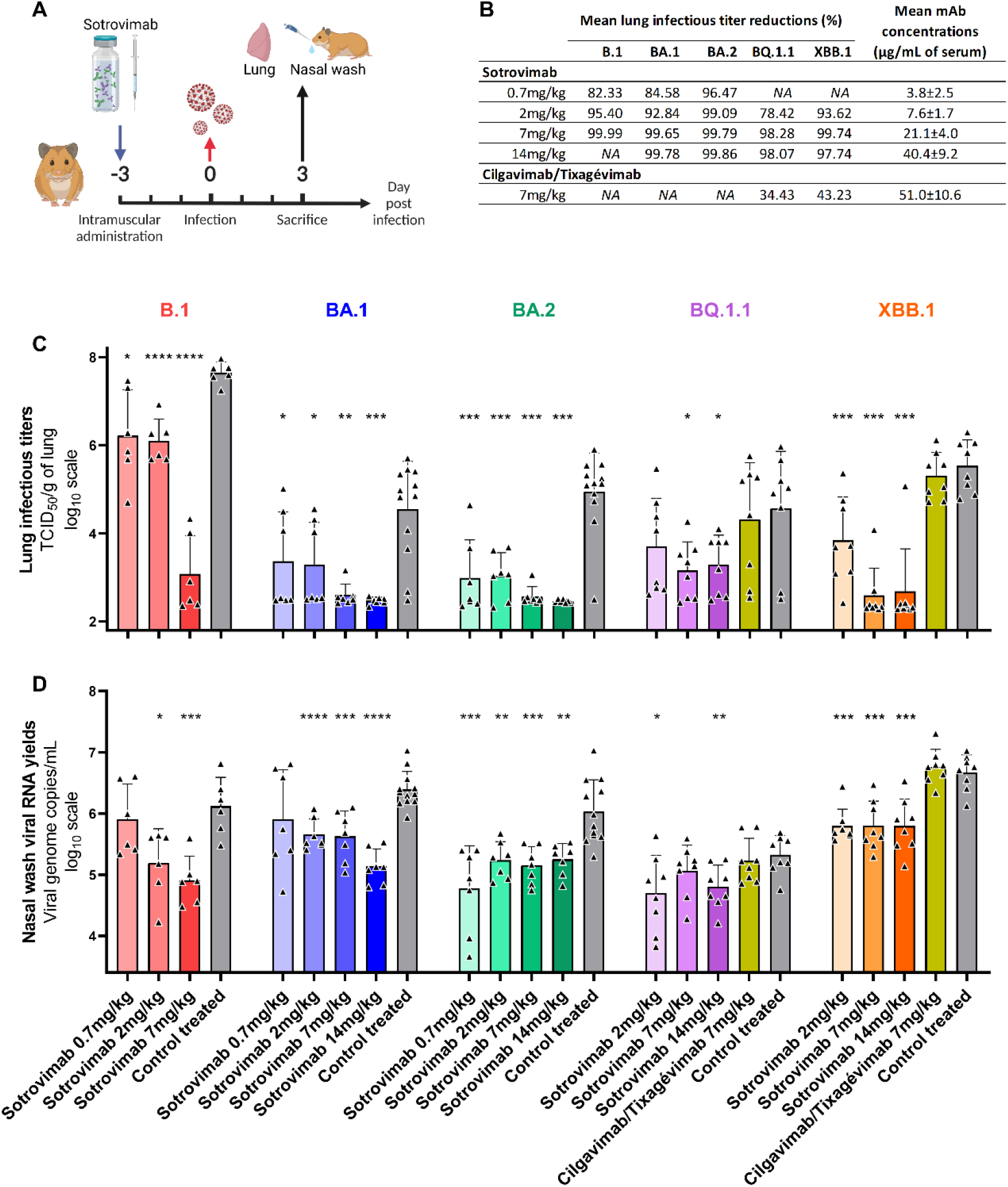
In vivo efficacy of Sotrovimab and Cilgavimab/Tixagévimab against BA.1, BA.2, BQ.1.1 and XBB.1 Omicron variants. (**A**) Experimental timeline. (**B**) Mean lung reduction infectious titer reduction compared to control treated animals and mean serum concentrations of mAbs at 3dpi. **(C**) Lung infectious titers and (**D**) viral RNA yields in nasal washes at 3dpi (Data represent mean ± SD of individual data). ****, ***, ** and * symbols indicate that the average value for the group is significantly lower than that of the control treated group with a p-value < 0.0001, ranging between 0.0001–0.001, 0.001–0.01, and 0.01–0.05, respectively.

Mean serum concentrations of Sotrovimab at 3dpi were measured and ranged between 3.8 and 40.4μg/mL (Fig 1B and S2). Pharmacokinetic data published by the US Food Drug Administration in humans indicate that mean serum concentrations peak at 143μg/mL and remain above 40μg/mL twenty-nine days after intravenous administration of 500mg Sotrovimab (COMET-ICE trial (NCT04545060)). These exposures are higher than those measured in groups of animals in which antiviral activity against the BQ1.1 and the XBB.1 variants is obtained, suggesting that Sotrovimab may retain some activity against these variants in treated patients.

Finally, a group of animals was also treated with 7mg/kg of Cilgavimab/Tixagévimab and infected with BQ.1.1 or XBB.1. No antiviral activity was obtained at this dose (Fig 1), in accordance with in vitro data published recently, and despite mean serum mAb concentrations above the geometric mean of 37.2μg/mL observed twenty-nine days after the administration of 600mg of this mAb cocktail during the TACKLE trial (NCT04723394) (Fig 1B and S2B).

Our data demonstrate that Sotrovimab retains some antiviral activity in vivo against the BQ.1.1 variant, although at a lower level than that observed against the first globally dominant BA.1 and BA.2 Omicron sublineages, in agreement with recently published in vitro data (Touret et al., 2022). In addition, treatment with S309 (parent antibody of Sotrovimab; 10 or 30 mg/kg) in mice or with Sotrovimab (10 mg/kg) in non-human primates also conferred protection against challenge with BQ.1.1 and corroborated our results (Addetia et al., 2023; Hérate et al., 2023). Regarding the XBB.1 variant, in our model, sotrovimab maintains a significant in vivo activity, comparable to that observed against the BA.1 and BA.2 Omicron sublineages. These results are in agreement with the activities observed in vitro using infected VeroE6/TMPRSS2 cells (Touret et al., 2022). While comparing exposure data in animals with those observed in humans, these findings support the potential efficacy of Sotrovimab against BQ.1.1 and underscore the urgent need to evaluate this mAb in humans. The clinical impact of Sotrovimab treatment against XBB.1 will need to be documented in more relevant models such as non-human primates and in humans. The constant antigenic evolution of SARS-CoV-2 reinforces the need for therapeutic antibodies and antiviral molecules with broad-spectrum activity and that may be used alone or in combination.

## Acknowledgments and funding

We thank Michela Scaffidi and Gregory Moureau for their technical contribution. We thank Pr. S. Van der Werf, Pr. B. Lina and Pr. C. Drosten for providing the SARS-CoV-2 strains. This work was performed in the framework of the Preclinical Study Group of the Emerging Infectious Diseases, the French research agency (ANRS-MIE). It was supported by the ANRS-MIE (BIOVAR and PRI projects of the EMERGEN research program) and by the European Commission (European Virus Archive Global project (EVA GLOBAL, grant agreement No 871029) of the Horizon 2020 research and innovation program). The funders of the study had no role in study design, data collection, data analysis, data interpretation, or writing of the report.

## Author contribution

Conceptualisation: JSD, XDL, AN

Data curation, formal analysis, investigation: JSD, OB, FT, AN

Funding acquisition, supervision: XDL, AN

Validation, writing – original draft: JSD, AN

Writing – review & editing: OB, FT, XDL

## Inclusion and diversity

We support inclusive, diverse and equitable conduct of research.

## Methods

### Cell line

VeroE6/TMPRSS2 cells were cultured at 37°C with 5% CO2 in minimal essential medium (MEM) supplemented with 1% Penicillin/Streptomycin, 1% non-essential amino acids, 7% of heat-inactivated fetal bovine serum (FBS) and 2% of G-418 disulfate 50mg/mL (all from ThermoFisher Scientific).

### Antibodies

Sotrovimab (Xevudy), Palivizumab (Synagis) and Cilgavimab/Tixagévimab (Evusheld) were obtained from pharmacy of the University hospital of La Timone, Marseille, France.

### Virus isolates

SARS-CoV-2 **B.1** strain BavPat1 was obtained from Pr. C Drosten through EVA GLOBAL (https://www.european-virus-archive.com/) and contains the D614G mutation. SARS-CoV-2 **Omicron BA.1** (B.1.1.529) was isolated the 1^st^ of December 2021 in Marseille, France. The full genome sequence has been deposited on GISAID: EPI_ISL_7899754. The strain, called 2021/FR/1514, is available through EVA GLOBAL (www.european-virus-archive.com, ref: 001V-04436). SARS-CoV-2 **Omicron BA.2** strain hCoV-19/France/NAQ-HCL022005338701/2022 was obtained from Pr. B Lina and the sequence is available on GISAID: EPI_ISL_9426119. SARS-CoV-2 **Omicron BQ.1.1** (hCoV-19/France/IDF-IPP50823/2022) was isolated by the National Reference Center for Respiratory Viruses hosted by Institut Pasteur (Paris, France) and headed by Pr. Sylvie van der Werf, from a specimen provided by Dr Beate Heym, Laboratoire des Centres de Santé et d’Hôpitaux d’IDF, 75020 Paris, France. The sequence is available on GISAID: EPI_ISL_15195982. SARS-CoV-2 **Omicron XBB.1.1** (hCoV-19/France/PAC-HCL022171892001/2022) was obtained from Pr. B Lina and the sequence is available on GISAID: EPI_ISL_15619797. All viral stocks were made by propagation in Vero E6/TMPRSS2 cells. All experiments with infectious viruses were performed in a biosafety level 3 laboratory.

### Study design

In vivo experiments were approved by the local ethical committee (C2EA—14) and the French ‘Ministère de l’Enseignement Supérieur, de la Recherche et de l’Innovation’ (APAFIS#35014). Three-week-old female Syrian hamsters (*Mesocricetus auratus*) were provided by Janvier Labs. Animals were maintained in ISOcage P-Bioexclusion System (Techniplast) with unlimited access to water/food and 14h/10h light/dark cycle. Animals were weighed and monitored daily for the duration of the study to detect the appearance of any clinical signs of illness, suffering or distress. Intramuscular administrations (in the hindlimb), infections, nasal washes and euthanasia (cervical dislocation) were performed under general anesthesia (isoflurane, Isoflurin^®^, Axience).

Group size was calculated with an effect size of 2 and a power of 80%, resulting in 6 to 12 animals/group. For untreated groups of hamsters (receiving an isotype control mAb): n=6 for the B.1 strain, n=12 for BA.1 and BA.2 variants, and n=8 for the BQ.1.1 and XBB.1.1 variants. For groups treated by Sotrovimab: n=6 for the B.1 strain, n=7 for BA.1 and BA.2 variants, and n=8 for the BQ.1.1 and XBB.1.1 variants. For groups treated by Sotrovimab and sacrificed before infection: n=6. For groups treated by Cilgavimab/Tixagévimab: n=8 for the BQ.1.1 and XBB.1.1 variants. Sample sizes were maximized within the capacity of the BSL3 housing and virus stock availability. Animals were randomly assigned to groups, but confounders were not controlled. Since, the same experimenters carried out infection/treatment/clinical follow-up, it was impossible to perform a blind trial. Predefined humane endpoints (>20% weight loss, moribund and a scoring >10 calculated according to a clinical evaluation scale) were set as exclusion criteria. No animals were excluded from the study.

### Experiment conduction

Four-to six-week-old anesthetized animals were intramuscularly inoculated with either a solution of Sotrovimab (0.7, 2, 7 or 14mg/kg), Cilgavimab/Tixagévimab (7mg/kg) or Palivizumab (7mg/kg; an isotype control mAb for ‘control treated’ animals) diluted in 0.9% sodium chloride. Three days later, they were intranasally infected with 50μL containing 1×10^4^ (B.1), 1×10^5^ (BA.1 and BA.2), 2×10^5^ (BQ.1.1) and 3×10^4^ (XBB.1.1) TCID50 of virus in 0.9% sodium chloride solution. To monitor exposure during the entire period in which the animals are infected, three additional groups of 6 uninfected animals were sacrificed three days after the intramuscular administration of Sotrovimab (2, 7 or 14mg/kg).

Nasal washes were performed at 3dpi: 150μl of 0.9% sodium chloride solution were instilled in the nasal cavities and transferred into 1.5mL tubes containing 0.5mL of 0.9% sodium chloride solution. Tubes were then centrifuged at 16,200g for 10 minutes and stored at −80°C. The animals were euthanised immediately following the completion of the nasal washes at 3 dpi. Lung samples were collected immediately after euthanasia. Left pulmonary lobes were washed with 10mL of 0.9% sodium chloride solution, blotted with filter paper, weighed, transferred into 2mL tubes containing 1mL of 0.9% sodium chloride solution and 3mm glass beads, crushed using a Tissue Lyser machine (Retsch MM400) for 20min at 30 cycles/s and centrifuged 10 min at 16,200g. Supernatant media were transferred into 1.5mL tubes, centrifuged 10 min at 16,200g and stored at −80°C. Two milliliters of blood were harvested in a 5mL serum tube. Blood was centrifuged for 10 min at 3,500g. Sera were transferred into 1.5mL tubes and stored at −20°C.

### Tissue-culture infectious dose 50 (TCID_50_) assay

96-well culture plates containing confluent VeroE6/TMPRSS2 cells were inoculated with 150μL per well of four-fold serial dilutions of each lung clarified homogenate samples. Each dilution was performed in sextuplicate. After 5 days of incubation, plates were read for the absence or presence of cytopathic effect in each well. Infectious titers were estimated using the method characterized by Reed & Muench (Reed, 1938).

### Quantitative real-time RT-PCR (RT-qPCR) assays

Nucleic acid from 100μL of nasal wash (spiked with bacteriophage MS2 (internal control)) were extracted using QIAamp 96 DNA kit and Qiacube HT robot (both from Qiagen). MS2 viral genome detection was performed as previously described(Driouich et al., 2021). Viral RNA yields were measured using a real time RT-qPCR assay targeting the RdRp coding region as previously described (Driouch et al., 2022; Driouich et al., 2021; Touret et al., 2021): Briefly, viral RNA was quantified by real-time RT-qPCR (GoTaq 1 step RT-qPCR kit, Promega). Quantification was provided by serial dilutions of an appropriate RNA standard. RT-qPCR reactions were performed on QuantStudio 12K Flex Real-Time PCR System (Applied Biosystems) and analyzed using QuantStudio 12K Flex Applied Biosystems software v1.2.3.

### Monoclonal antibodies quantification

To estimate the exposure of animals to monoclonal antibodies, we measured the level of human serum IgG antibodies directed against the S1 domain of the spike protein of the SARS-CoV-2 using a commercial enzyme-linked immunosorbent assay (ELISA) kit (Euroimmun) following manufacturer instructions. Results were expressed in binding antibody units per mL (BAU/mL) following manufacturer instructions and converted to μg/mL using blank plasma from untreated/infected animals spiked with known quantities of monoclonal antibodies.

### Interpretation of the results

Graphical representations and statistical analyses were realized using Graphpad Prism 7 software. Two-sided statistical analysis were performed using Shapiro–Wilk normality test, Fisher’s exact test, Student t-test, Mann–Whitney test and Welch’s test. P-values lower than 0.05 were considered statistically significant. Lung infectious titers reductions were estimated as follows: lung infectious titer of an animal divided by the mean titer of the untreated group infected with the corresponding virus. Experimental timelines (Figure 1A) were generated on biorender.com

---

**Figure S1:**
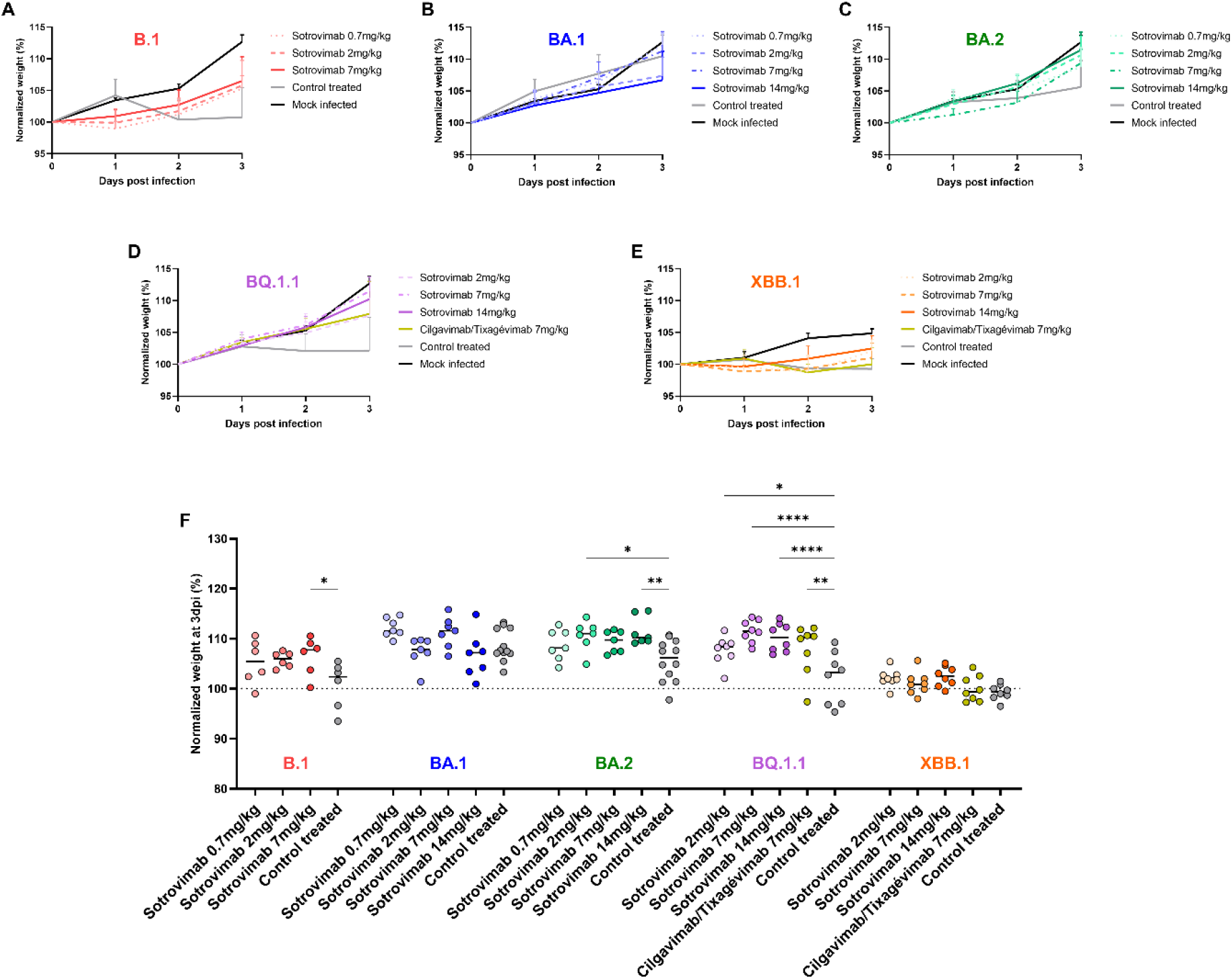
Clinical monitoring of animals. **(A-E**) Normalized weights (% of initial weight of the animal) of animal infected by the ancestral B.1 strain, and the Omicron BA.1, BA.2, BQ.1.1 or XBB.1 variants. (**F**) Individual normalized weights at 3dpi of animal infected by the ancestral B.1 strain, the Omicron BA.1, BA.2, BQ.1.1 or XBB.1 variants. ****, ** and * symbols indicate that the average value for the group is significantly different than that of the control treated group with a p-value < 0.0001, ranging between 0.001–0.01, and 0.01–0.05, respectively.

**Figure S2:**
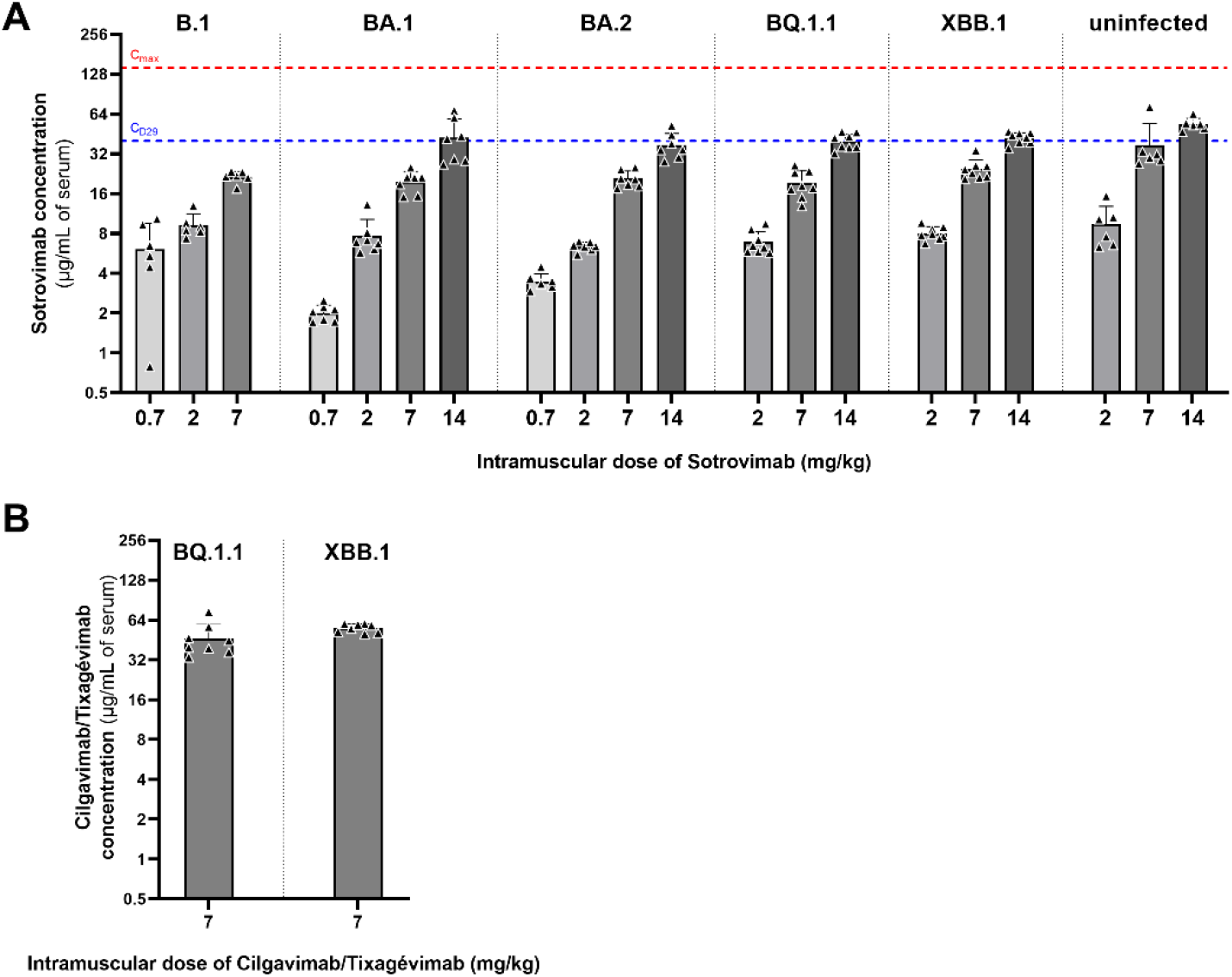
Serum mAb concentrations. (**A**) Serum concentrations of Sotrovimab in infected animals sacrificed at 3dpi, and in uninfected animals receiving the mAb three days before. Red and blue dotted lines indicate respectively maximum (C_max_) and day 29 (CD_29_) mean serum concentrations observed after intravenous administration of 500mg of Sotrovimab in humans (data published by the US Food Drug Administration; COMET-ICE trial (NCT04545060); https://www.fda.gov). (**B**) Serum concentrations of Cilgavimab/Tixagévimab in infected animals sacrificed at 3dpi.

